# AURKA destruction is decoupled from its activity at mitotic exit but suppresses interphase activity

**DOI:** 10.1101/850917

**Authors:** Ahmed Abdelbaki, H. Begum Akman, Marion Poteau, Rhys Grant, Olivier Gavet, Giulia Guarguaglini, Catherine Lindon

## Abstract

Activity of AURKA is controlled through multiple mechanisms including phosphorylation, ubiquitin-mediated degradation, and allosteric interaction with TPX2. Activity peaks at mitosis before AURKA is degraded during mitotic exit in a process strictly dependent on APC/C coactivator FZR1. We used FZR1 knockout cells (FZR1^KO^) and a novel FRET-based AURKA biosensor to investigate how activity is regulated in absence of destruction. We found that AURKA activity in FZR1^KO^ cells dropped at mitotic exit as rapidly as in parental cells, despite absence of destruction. Unexpectedly, TPX2 was degraded normally in FZR1^KO^ cells. Overexpression of an N-terminal TPX2 fragment sufficient for AURKA binding, but not degraded at mitotic exit, caused delay in AURKA inactivation. We conclude that AURKA inactivation in mitotic exit is determined not by its own degradation but by degradation of TPX2 and therefore dependent on CDC20 rather than FZR1. The biosensor revealed that FZR1 instead suppresses AURKA activity in interphase and is critically required for assembly of the interphase mitochondrial network after mitosis.

## Introduction

Aurora kinase A (AURKA) is a major mitotic kinase required for multiple steps in mitosis, including centrosome maturation, microtubule nucleation and organisation into a bipolar spindle and mitotic checkpoint function [1, 2]. Recent studies of the effects of acute inhibition of AURKA during mitosis have concluded that it plays an important role during mitotic exit in regulating assembly of the anaphase spindle upon which sister chromatids are segregated [3–5]. AURKA also has a number of non-mitotic roles that include disassembly of the primary cilium, regulation of myc family transcription factors and response to replication stress [6–10]. We and others recently reported that AURKA constitutively regulates mitochondrial morphology and function [11, 12] in addition to a previously reported role in promoting mitochondrial fission in mitosis [13].

Widespread reports in the literature of interphase activity of AURKA, and of its activation through a number of interacting partners [14], can be explained by the unique allosteric properties of this kinase. Either autophosphorylation of the T-loop [15–17] or allosteric interactors [18–21] contribute to the active conformation of AURKA. The most prominent of these allosteric activators is TPX2, which controls AURKA localization, activation and stability on the mitotic spindle during mitosis [16, 22, 23]. Interaction of TPX2, whilst protecting the autophosphorylated site (pT288 in hsAURKA) from access by PP1 phosphatase [16], also acts to stabilize the T-loop with or without its phosphorylation. Consequently, phosphatase-mediated reversal of T288 phosphorylation may not be sufficient to eliminate activity of AURKA.

In somatic cells, both AURKA protein and kinase activity are low or undetectable through much of the cell cycle, rising sharply in G2 phase, most prominently on the duplicated centrosomes and then on the bipolar spindle. pT288 staining is almost entirely confined to the centrosomes, in line with the idea that autophosphorylation is just one route to activation of AURKA. Indeed a conformational sensor of AURKA detects active kinase on the spindle, and in the cytoplasm, both during and after mitosis [24]. Mitotic destruction of AURKA begins late in anaphase following assembly of the spindle midzone and at the time of spindle pole disassembly, in a manner that depends on the Cdh1 co-activator (hereafter referred to as FZR1) of the Anaphase Promoting Complex/Cyclosome (APC/C) [25–28].

The APC/C is responsible for destruction of dozens of substrates during mitotic exit and coordinates mitotic exit with the processes of chromosome segregation and cytokinesis through the concerted regulation of its coactivators [29]. APC/C-CDC20 targets cyclin B and securin during metaphase to drive chromosome segregation and mitotic exit. After anaphase onset a switch in specificity of APC/C-CDC20 and/or activation of APC/C-FZR1 degrades all the remaining substrates including Aurora kinases. Like most APC/C substrates, AURKA destruction is efficiently accomplished by APC/C-FZR1 in a variety of different assays. Unlike most substrates, AURKA and AURKB show strict dependence on FZR1 for their destruction in cell-based assays [28, 30, 31]. Neither the molecular basis, nor the functional significance of this specificity in AURK targeting, is understood.

Given the complexity of AURKA activity during the cell cycle, we investigated what the destruction of AURKA contributes to its inactivation during mitotic exit using novel tools in the form of a CRISPR/Cas9 FZR1 knockout cell line we have recently generated and a FRET-based AURKA activity biosensor. We conclude from our studies that FZR1-mediated destruction of AURKA plays no role in the timing of its inactivation during mitotic exit but is critical to suppress interphase activity and function of the kinase. We test this idea by demonstrating that reestablishment of mitochondrial connectivity after mitosis requires suppression of activity of undegraded AURKA.

## Results

Although AURKA activity has been described to peak in M phase there has been no detailed characterization of how AURKA activity varies through mitosis. In particular the question of how AURKA activity is regulated during mitotic exit, given that AURKA also has anaphase functions [32, 33], and the importance of its ongoing destruction for attenuation of activity, remain unresolved. We first used a commercially available antibody against the activated T-loop of Aurora kinases (pT288 in AURKA) that confirms by immunoblot the peak of active AURKA at mitosis in extracts from synchronised U2OS cells (Figure 1A). We then used the same antibody to look by immunofluorescence (IF) in fixed mitotic cells, marking centrosomes with γ-tubulin staining. All phospho-epitope signal on the centrosomes and spindle poles was abolished by treatment with AURKA inhibitor MLN8237, whereas the signal at the midbody (where the same antibody recognizes pT232 of AURKB) was insensitive to 100 nM MLN8237, indicating that the centrosomal pT288 signal is specific to AURKA as expected (Figure 1B, Supplementary Figure S1) [34, 35]. We quantified fluorescence intensity associated with pT288 at different stages of mitosis scored according to DNA morphology. We found that pT288 was evident at the centrosomes from G2 phase onwards. Following nuclear envelope breakdown (NEB), prometaphase (PM) cells showed a strong increase in centrosome- and spindle pole- associated pT288 signal, peaking at metaphase. pT288 signal remained high at anaphase before starting to fall significantly in telophase cells (Figure 1C, D). We concluded that inactivation of AURKA through dephosphorylation of its T-loop occurs at around the time of onset of AURKA destruction, which occurs about 10 minutes after anaphase onset in human cells [28]. Immunoblot analysis of extracts from cells synchronized through mitotic exit using a spindle assembly checkpoint (SAC) arrest/release protocol confirmed that pT288 levels and total AURKA both fall rapidly during forced mitotic exit (Figure 1E).

**Figure 1:**
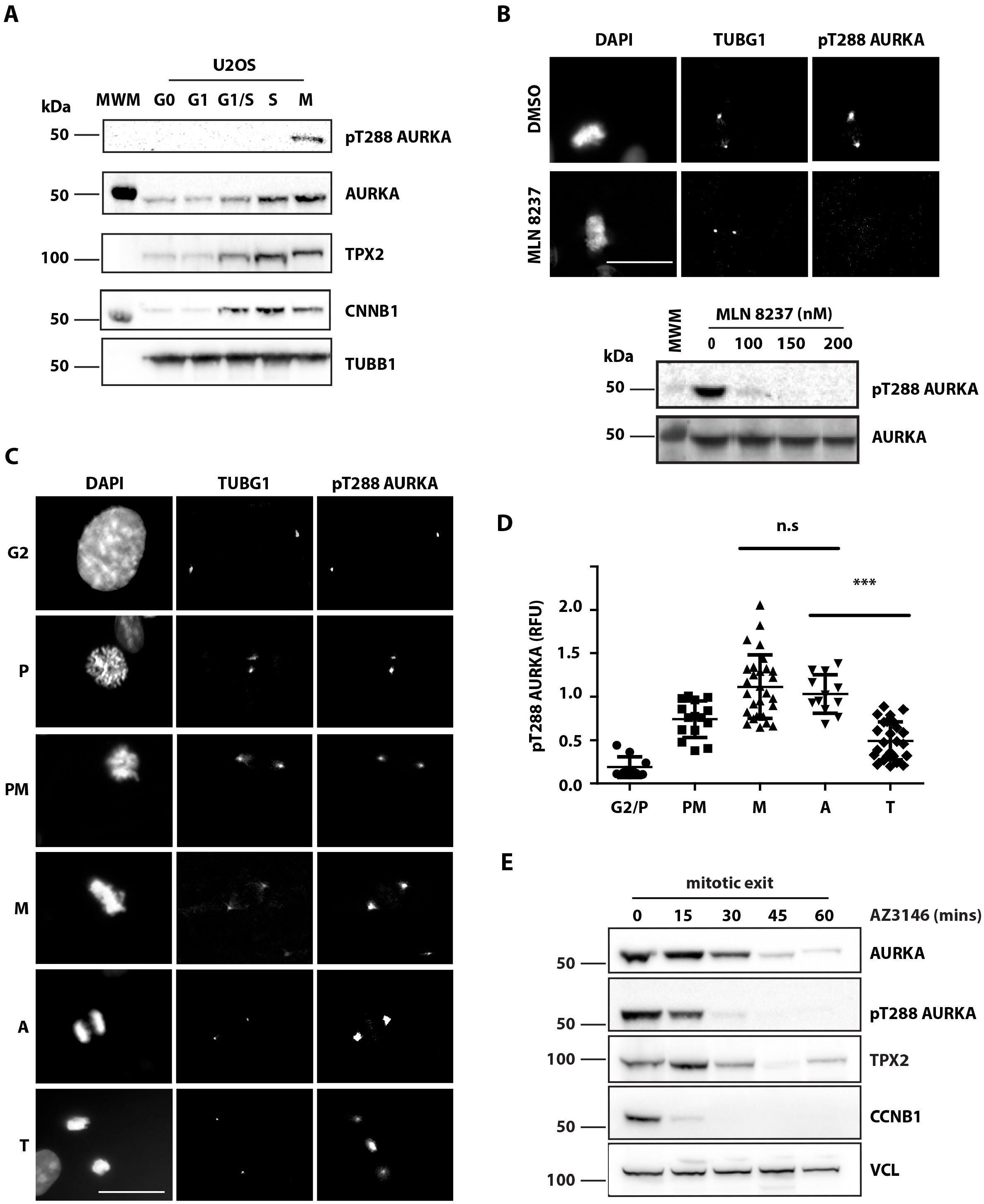
Inactivation of AURKA during mitotic exit begins at anaphase. **A** pT288 antibody detects active AURKA only in mitotic cells. Cells were synchronized as described in Materials and Methods and blotted for pT288, total AURKA and other mitotic markers. **B** pT288 signal is sensitive to AURKA-specific inhibitor MLN8237 by IF on mitotic cells from a MeOH-fixed unsynchronized population (upper panel) or by immunoblot of STLC-arrested mitotic cells treated for 3 hours at the indicated doses (lower panel). AURKA-specific pT288 signal is restricted to centrosomes and spindle pole bodies (marked by γ-tubulin, TUBG1). Bars, 10 μm. See also Figure S1. **C-E** Quantification of pT288-AURKA during mitotic exit. **C, D** Unsynchronized cell populations were fixed and stained as in **B**. Cells were judged to be at different stages of mitosis according to DAPI staining (**C**) and scored for mean pT288 AURKA signal measured in a fixed ROI centred on TUBG1 signal at centrosomes or spindle poles (**D**). G2 and prophase (P), n=10; prometaphase (PM), n=15; metaphase (M), n=30; anaphase (A), n=30; and telophase (T), n=26. M vs A, not significant (n.s.); A vs T, p < 0.0001 (***), Students’ t-test. **E** Cells were synchronized in 5 μM STLC and released by checkpoint inhibition using 10 μM AZ3146, with extracts harvested at times indicated. These were examined by immunoblotting for AURKA, pT288-AURKA and TPX2 levels. Disappearance of Cyclin B1 (CCNB1) acts as marker for mitotic exit, level of vinculin (VCL) as loading control.

Since the kinetics of dephosphorylation and destruction of AURKA in mitotic exit were not identical (Figure 1E), we investigated whether AURKA destruction contributes to the fall in kinase activity at mitotic exit. Mitotic AURKA destruction is critically dependent on the FZR1 co-activator of APC/C, so we used a FZR1 knockout (FZR1^KO^) in U2OS cells generated by CRISPR/Cas9 (Supplementary Figure S2) to study AURKA inactivation in the absence of its destruction at mitotic exit. In FZR1^KO^ cells, AURKA-Venus protein levels measured in single cell degradation assays remained constant through mitotic exit, compared to the parental U2OS cell line (Figure 2A), and as previously reported using siRNA-mediated suppression of FZR1 [28]. We then compared loss of pT288 staining in mitotic exit between parental and FZR1^KO^ cells using the assays established in Figure 1. We found by immunoblot that loss of pT288 signal was identical in both cell lines during synchronized mitotic exit, despite strong stabilization of the AURKA signal in FZR1^KO^ cells (Figure 2B). AURKB was also stabilized but less so, consistent with slower degradation of AURKB [36]. We note that degradation of endogenous TPX2 during mitotic exit, like inactivation of AURKA, was insensitive to FZR1 knockout, and appeared more complete in FZR1^KO^ cells, where CDC20 levels persisted compared to the parental cells (Figure 2B). TPX2 was first proposed as a substrate of the Cdh1/FZR1-activated version of APC/C [37] but our results suggest that in an unperturbed mitotic exit TPX2 – like cyclin B1 (CCNB1) – is a substrate for CDC20 ahead of FZR1 activation. We then fixed parental U2OS cells and FZR1^KO^ cells and processed them for quantitative IF analysis of pT288-AURKA and total AURKA staining at spindle poles at metaphase and telophase. In this analysis AURKA persisted on spindle poles during mitotic exit in FZR1^KO^ cells compared to parental U2OS, but with no corresponding persistence in pT288 label (Figure 2C). In both cell lines, the presence of active AURKA, as measured by T-loop phosphorylation of the kinase, was strongly reduced after onset of mitotic exit independent of the level of protein remaining. We confirmed that the persistence of AURKA on spindle poles is most likely a direct effect of AURKA non-degradation, since wild-type AURKA-Venus, but not a non-degradable version, is rapidly lost from spindle poles in unperturbed mitotic exit (Supplementary Figure S2).

**Figure 2:**
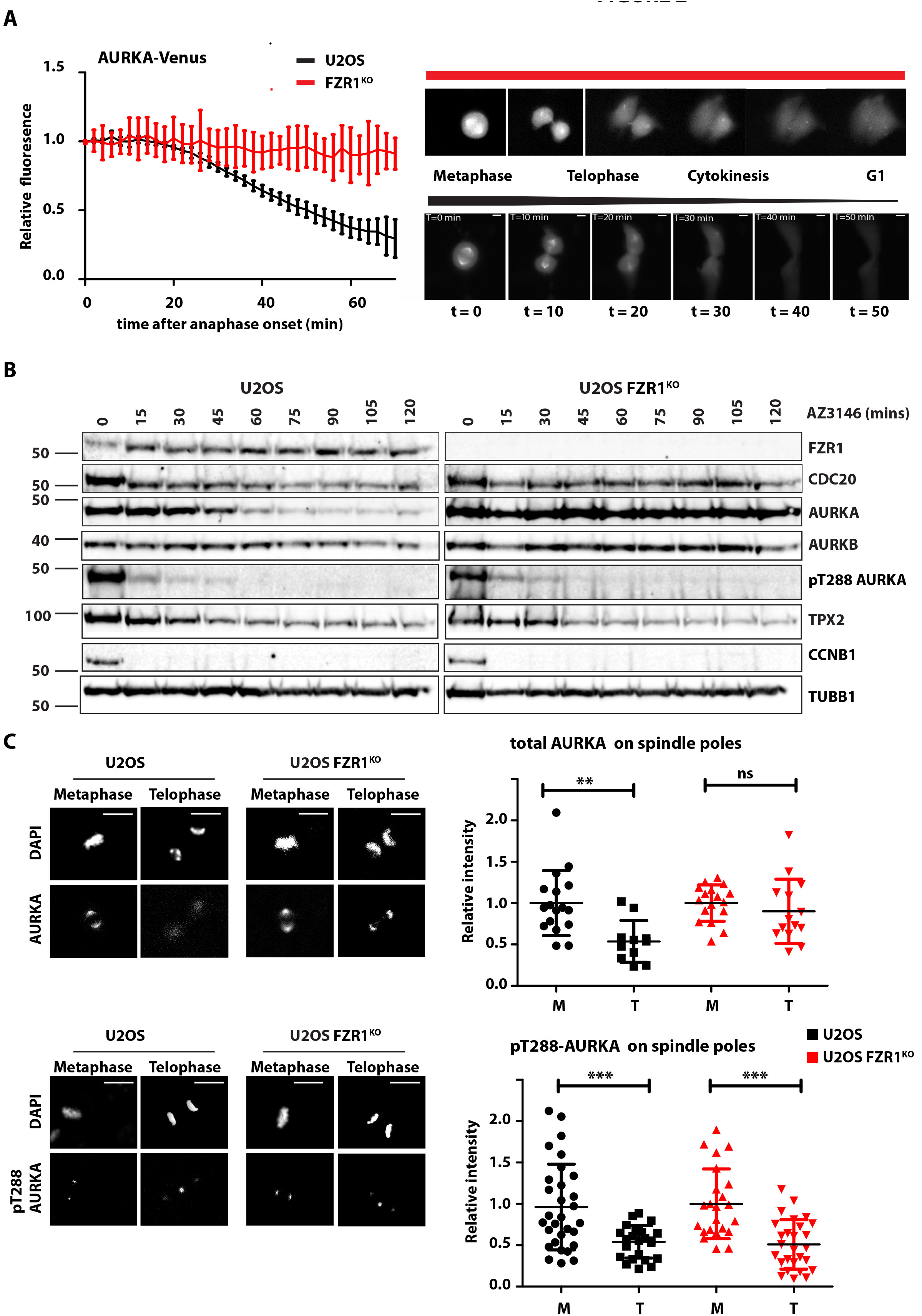
AURKA destruction is not required for pT288-AURKA down-regulation at mitotic exit. **A** There is no mitotic exit destruction of AURKA in FZR1/Cdh1 KO cells (FZR1^KO^). AURKA-Venus was transiently transfected into both U2OS and U2OS FZR1^KO^ cells. Quantifications of total fluorescence measurements from single mitotic cells (illustrated in righthand panels) were used to generate degradation curves for AURKA-Venus (lefthand panel). Fluorescence values for individual curves were normalized to the last frame before anaphase onset, and all curves *in silico* synchronized to anaphase. n = 10 cells. **B-D** AURKA activity scored by pT288 is not affected by FZR1^KO^ during mitotic exit. **B** U2OS and FZR1^KO^ cells were synchronized to prometaphase using 5 μM STLC and released by checkpoint inhibition using 10 μM AZ3146, with extracts harvested at times indicated. Lysates were analyzed by immunblot with antibodies against AURKA, pT288-AURKA and other mitotic regulators. **C, D** pT288-AURKA and AURKA staining associated with individual centrosomes/spindle poles in metaphase (M) versus telophase (T) cells (lefthand panels). Fluorescence values were measured as in Figure 1 and are presented as dot plots for total AURKA (**C**) and pT288-AURKA (**D**) in both U2OS and FZR1^KO^. All values were normalized to the mean value from control metaphase cells. ns, non-significant; ** p < 0.001; *** p < 0.0001, Student’s t-test. **C**, n ≥ 11; **D**, n ≥ 23. Bars, 10 μm.

A number of AURKA binding partners have been shown to affect the activity of AURKA, some independently of phosphorylation on T288, via distinct effects on its conformational dynamics. Therefore, we used a diffusible kinase biosensor, based on the well-established design of a fluorescent protein FRET pair separated by a phospho-threonine binding domain and specific phosphorylation motif [38], to provide a more comprehensive readout of AURKA activity. An AURKA-directed version of this biosensor was created using the T210 motif from the T-loop of Plk1 as a known target of AURKA activity [39, 40]. Phosphorylation of the T210 motif in the biosensor causes loss of FRET in mitotic cells, measured as an increase in CFP/YFP emission ratio of approximately 10%, in a manner dependent on its phosphorylation site and sensitive to specific inhibitors of AURKA (Figure 3A-B, Supplementary Figure S3). Some sensitivity to inhibitors of AURKB at higher doses indicated that the biosensor might not be completely specific to AURKA. (Supplementary Figure S3). This finding was not unexpected for a diffusible biosensor, since some of the specificity in substrate phosphorylation by Aurora kinases is proposed to reside in the colocalization of kinase with substrates [3, 35]. We transfected this biosensor into U2OS and FZR1^KO^ cells to measure Aurora kinase activity at mitosis and during mitotic exit. FRET measurements in mitotic cells were in agreement with the analysis of pT288 staining shown in Figure 2, in showing that biosensor activity started to increase in late G2, peaking after NEB and decaying during mitotic exit. We found that the increase in activity measured at mitotic entry was identical in individual U2OS and FZR1^KO^ cells *in silico* synchronized to NEB (Figure 3D): peak activity showed a small but not significant increase in FZR1^KO^ cells. *In silico* synchronization to anaphase onset (Supplementary Figure S3) revealed that the timing of Aurora kinase inactivation was identical. Moreover, if we normalized FRET signals to the anaphase onset value, the inactivation curves were directly superimposable (Figure 3E). Therefore, mitotic Aurora kinase inactivation is independent of FZR1. FRET measurements returned to pre-mitotic levels at about 1 hour after anaphase onset, indicating complete inactivation of the mitotic pool of Aurora kinases. We observed, however, that biosensor activity starts to increase again gradually in G1 in FZR1^KO^ cells compared to parental U2OS cells and is significantly increased at 160 minutes after NEB (Figure 3D). We concluded that destruction of AURKA does not influence its inactivation during mitotic exit but may be important to prevent re-activation early in the cell cycle.

**Figure 3:**
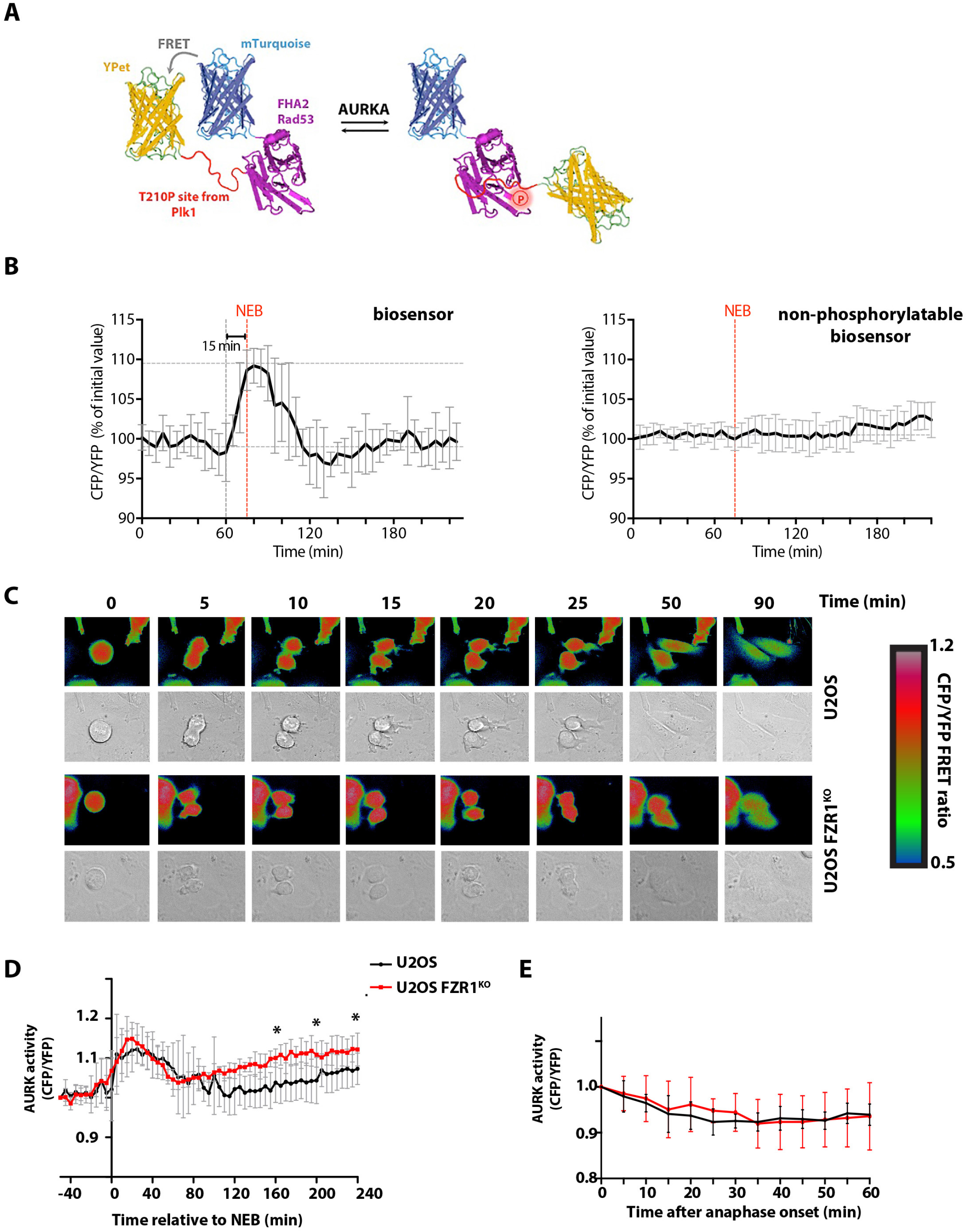
A FRET-based biosensor records unaltered parameters of mitotic AURKA activation and inactivation in FZR1^KO^ cells. **A** Schematic illustration of AURKA biosensor showing high FRET (left) versus low FRET states (right). **B** Inverted FRET measurements (CFP/YFP emission) from timelapse movies of cells expressing the biosensor, or a non-phosphorylatable version, show that the biosensor reports on mitotic phosphorylation events, n ≥ 8. Further characterization of the specificity of the biosensor is shown in Supplementary Figure S3. **C** Examples of inverted false-coloured FRET ratio of biosensor-expressing single U2OS and FZR1^KO^ cells passing through mitosis: High FRET (blue) reports on non-phosphorylated state, whereas low FRET (red) reports on the phosphorylated probe. **D-E** FRET ratio values measured using biosensor show AURK activity is normally regulated through mitosis in FZR1^KO^ cells but rises again in G1 phase (*, p < 0.05, Student’s t-test). **D** Cells *in silico* synchronized to NEB. **E** Cells *in silico* synchronized and FRET values normalized to anaphase onset.

How can we explain the timing of AURKA inactivation at mitotic exit? We have previously shown that interaction with TPX2 stabilizes AURKA against APC/C-FZR1-mediated degradation [23], and from this concluded that loss of interaction with TPX2 contributes to the timing of AURKA degradation in mitotic exit. Our finding here that AURKA degradation plays no role in its inactivation led us to evaluate whether AURKA inactivation also could depend on TPX2 degradation. Since transient overexpression of full-length TPX2 inhibited mitotic progression in our hands, we used an N-terminal fragment of TPX2 (amino acids 1-43) known to be sufficient for binding to and activating AURKA [16]. We overexpressed TPX2(1-43)-CFP in U2OS and FZR1^KO^ cells and monitored pT288-AURKA during mitotic exit (Figure 4). We found that persistence of TPX2(1-43) through mitotic exit stabilized AURKA protein during mitotic exit in U2OS cells (Figure 4A,B) as previously described [23]. As expected, TPX2(1-43) showed no effect on AURKA levels in FZR1^KO^ cells where the protein is not degraded (Figure 4C,D). Importantly, we found a marked effect of TPX2(1-43) in stabilizing the pT288-AURKA signal during mitotic exit in both parental U2OS cells and FZR1^KO^ cells (Figure 4A-D). We concluded that loss of interaction with TPX2 is the rate-limiting step in inactivation of AURKA in mitotic exit. We confirmed that inhibition of the candidate phosphatase, PP1, after mitotic exit also stabilized pT288 signal (Figure 4E), consistent with the conclusion that de-activation rather than destruction controls AURKA activity in mitotic exit.

**Figure 4:**
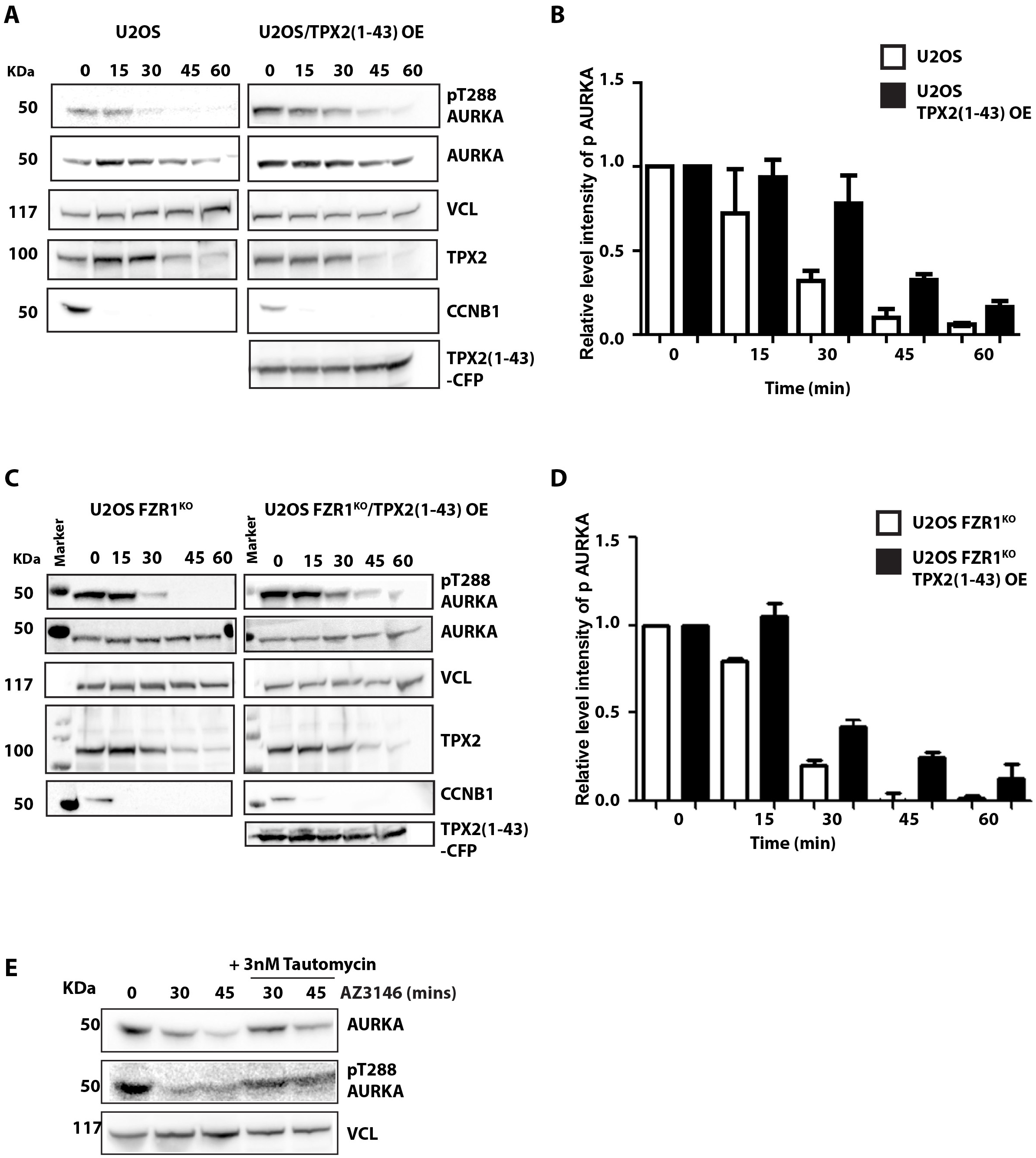
AURKA inactivation is controlled through TPX2. U2OS (**A,B**) and FZR1^KO^ (**C,D**) cells were transfected with TPX2(1-43)-CFP and synchronized through mitotic exit as described in the legend to Figure 2. Quantitative immunoblotting of cell lysates shows that loss of pT288-AURKA during mitotic exit is delayed in the presence of TPX2(1-43) in both parental and FZR1^KO^ cells. Cyclin B1 (CCNB1) is used as marker for mitotic exit, level of vinculin (VCL) as loading control. Bar charts **(B, D)** show pT288 signal normalized against vinculin. Results presented are mean values from 3 independent experiments ± S.D. **E** AURKA inactivation is phosphatase dependent. U2OS cells undergoing mitotic exit were treated with PP1 inhibitor 3nM tautomycin 10 minutes after relief of checkpoint inhibition by AZ3146. Lysates harvested at the indicated time points after AZ3146 treatment were subject to immunoblot analysis.

If FZR1-mediated destruction of AURKA is not required for its inactivation at mitotic exit, then what is it for? We noticed a small but significant increase in accumulation of Aurora kinase biosensor activity following mitotic exit in FZR1^KO^ cells (Figure 3D). Using both AURKA and AURKB biosensors [41] and pT288 staining, we examined Aurora kinase activity in cells arrested at the G1/S boundary by double thymidine block. The AURKA biosensor revealed increased Aurora kinase activity in FZR1^KO^ cells that was abolished by treatment with MLN8237 (Figure 5A), but insensitive to a low dose of AZD1152 inhibitor sufficient to abolish AURKB-specific activity (Supplementary Figure S3). Since the AURKB sensor is insensitive to MLN8237 at the dose used (Supplementary Figure S4), we concluded that there is increased AURKA activity in interphase in FZR1^KO^ cells. Consistent with this conclusion, we found that pT288 stained centrosomes in interphase FZR1^KO^ cells but not parental U2OS (Figure 5B).

**Figure 5:**
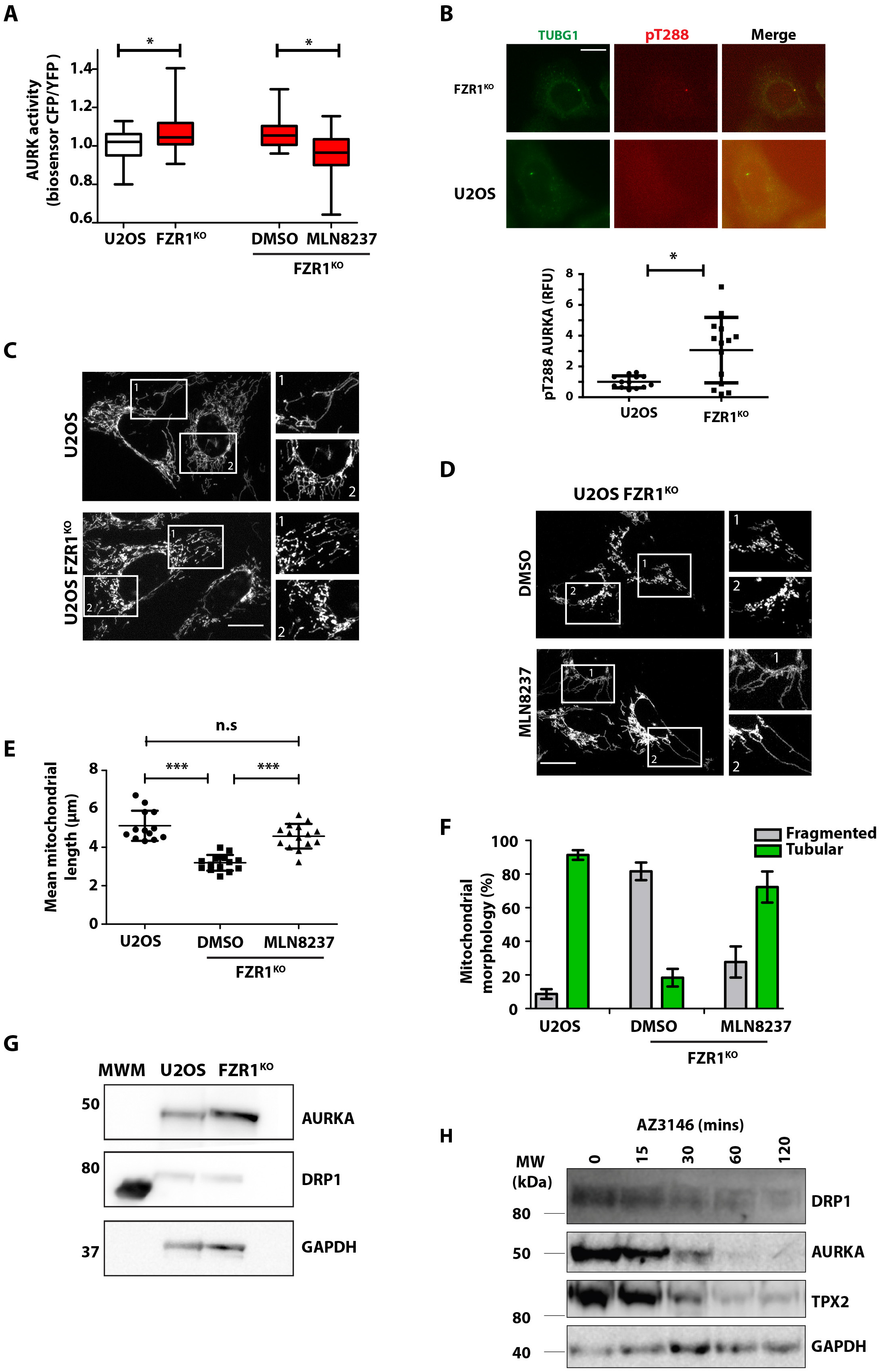
Destruction of AURKA by APC/C^Cdh1/FZR1^ is required to control interphase activity of AURKA. **A** The FRET biosensor reveals raised Aurora kinase activity in interphase FZR1^KO^ cells. Cells were synchronized in G1/S using double thymidine block. Box and whisker plots show CFP/YFP emission ratios measured for U2OS and FZR1^KO^ cell populations (n = 20; p < 0.05, Student’s t-test). The increased activity measured in FZR1^KO^ cells was sensitive to treatment with AURKA inhibitor MLN8237, 100nM for 3 hours. **B** pT288 staining could be detected at the centrosomes of G1/S FZR1^KO^ cells, but not parental U2OS cells, processed for IF analysis **C-F** Mitochondria are over-fragmented in interphase FZR1^KO^ cells in an AURKA-sensitive manner. U2OS and FZR1^KO^ cells were synchronized in G1/S then stained with MitoTracker^®^ and imaged live (**C**). FZR1^KO^ cells were treated with DMSO or 100nM MLN8237 for 3 hours (**D**). Scale bars, 10 μm. **E** Quantitative analyses of fragmented mitochondria length was carried out as described in Materials and Methods and are presented as dot plots (n ≥ 15 cells; p < 0.001 by 2-tailed Mann-Whitney U test), whilst (**F**) percentages of tubular versus fragmented morphologies were calculated using MicroP software and are presented as mean values ± S.D (n ≥ 15; p < 0.0001 by 2-tailed Mann-Whitney U test). **G-H** Immunoblotting U2OS and FZR1^KO^ cells shows that DRP1 levels, unlike those of AURKA, are not altered in G1/S FZR1^KO^ cells (**G**), even though DRP1 undergoes modest degradation during mitotic exit (**H**).

Next, we explored how FZR1 affects known AURKA functions in interphase. We and others have previously described a role for AURKA, at physiologically relevant levels of expression, in promoting mitochondrial fission during interphase [11, 12], an observation thought to be significant to cellular metabolism given the intimate link between mitochondrial morphology and function [42]. We hypothesized therefore that one consequence of increased interphase AURKA activity should be increased mitochondrial fragmentation in FZR1^KO^ cells, and indeed mitochondrial fragmentation following FZR1 siRNA treatment of HeLa cells has previously been described [43]. We compared the mitochondrial networks in parental U2OS and FZR1^KO^ cells arrested at the G1/S boundary, when the mitochondria tend to form large interconnected mitochondrial networks [44, 45], and found the mitochondrial network to be highly fragmented in FZR1^KO^ cells in a manner dependent on AURKA activity (Figure 5C-F). We measured reduced mean mitochondrial lengths in FZR1^KO^ cells (Figure 5E), and classified mitochondria according to morphological subtypes, to find a significant increase in the percentage of fragmented globules and significantly reduced tubular morphology in FZR1^KO^ cells (Figure 5F). Horn and co-authors had identified the fission factor dynamin-related protein-1 (DRP1) as the target of FZR1 required for its effect on mitochondrial morphology [43]. Here we found that inhibition of AURKA activity completely rescued mitochondrial length and morphology in FZR1^KO^ cells (Figure 5E,F), consistent with data showing AURKA to be an upstream regulator of DRP1 [11, 13]. Treatment of FZR1^KO^ cells with an AURKB-specific inhibitor had no effect on mitochondrial length (Supplementary Figure S4). Moreover we found no alteration in DRP1 levels in FZR1^KO^ cells (Figure 5G) although, in agreement with [43], we observed loss of DRP1 during mitotic exit in parental cells (Figure 5H) and suggest that it could be a substrate for APC/C-CDC20 as well as APC/C-FZR1, or another ubiquitin ligase. We concluded that APC/C-FZR1 regulates mitochondrial dynamics by preventing AURKA reactivation in interphase.

To further test our hypothesis that mitochondrial fragmentation is a direct response to un-degraded AURKA, we tested if the introduction of nondegradable (nd, Δ32-66) and wild-type (WT) versions of AURKA would show differential effects on mitochondrial dynamics (Figure 6). We used stable RPE1-FRT/TO cell lines expressing either AURKA-Venus-WT or AURKA-Venus-nd under tetracycline control. Induction of AURKA-Venus-WT caused increased fragmentation in unsynchronized cells as previously described [12], but AURKA-Venus-nd had a greater effect (Figure 6A,B). Importantly, this mitochondrial phenotype was independent of any effect of AURKA on microtubule dynamics that might contribute to subcellular distribution of mitochondria (Supplementary Figure S5). We then tracked individual cells through mitotic exit in the presence of MitoTracker^®^. Treatment with MLN8237 interfered with mitochondrial fragmentation at mitosis as expected from the published findings [13] (Figure 6C). In the presence of overexpressed AURKA-Venus we found that mitochondria were excessively fragmented as cells progressed out of mitosis into G1 phase, and that this effect was exacerbated in cells expressing AURKA-Venus-nd compared to WT (Figure 6D). We concluded that undegraded AURKA is able to inhibit reassembly of the interphase mitochondrial network, and that AURKA is a direct target of FZR1 in regulating mitochondrial dynamics.

**Figure 6:**
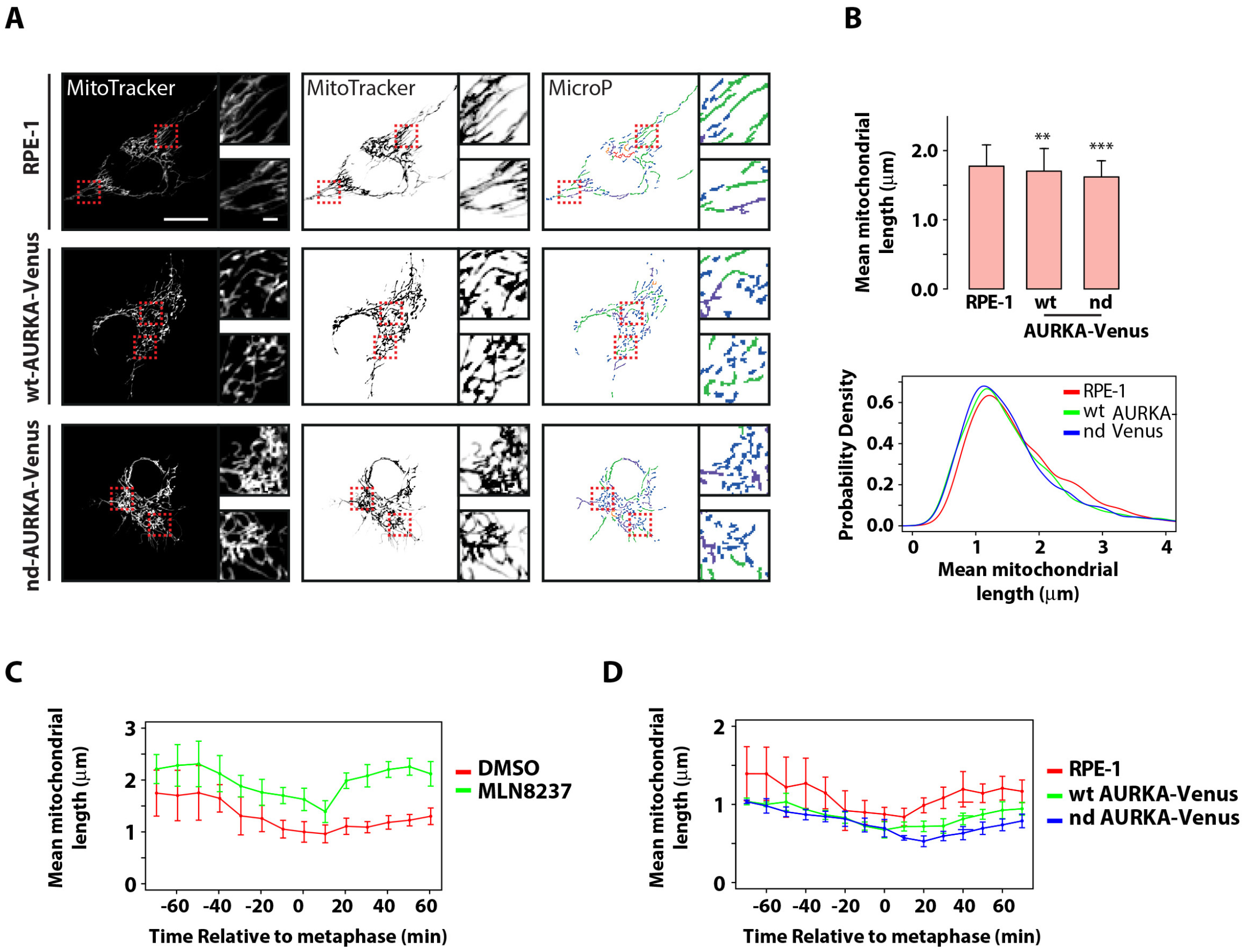
Undegraded AURKA at mitotic exit inhibits reassembly of the interphase mitochondrial network. **A,B** RPE-1, RPE-1 AURKA-Venus-WT and non-degradable version (nd, Δ32-66) cell lines were stained with MitoTracker^®^ and imaged 24 h after induction of AURKA transgene expression. Acquired images were analysed with MicroP(**A**) and subjected to mitochondria length quantifications (**B**). n = 30 mitochondria per cell, 18-24 cells per condition, two experimental replicates plotted as mean values ± S.D(upper panel) or as Kernel density plots (lower panel). **C** RPE1 cells treated with either DMSO or 100 nM MLN8237 were stained with MitoTracker^®^ and filmed live as they progressed through mitosis. Mean mitochondrial length over time is plotted for n ≥ 6 cells per condition in two experimental replicates. **D** RPE-1 AURKA-Venus-WT and -nd cells were stained with MitoTracker^®^, filmed and analysed as in **C**, n ≥ 5 cells.

## Discussion

AURKA activity increases in preparation for mitosis in parallel with the protein level, and both progressively drop from anaphase onwards. This has led to the expectation that destruction contributes to regulating AURKA activity at mitotic exit [28, 33]. We observed, however, that loss of AURKA activity proceeds faster than destruction of the protein. Lack of the APC/C co-activator FZR1 stabilizes AURKA levels but does not affect the timing of AURKA inactivation during mitotic exit, as measured using either pT288 reactivity or a novel FRET biosensor for AURKA activity that we describe in this study. Therefore, AURKA destruction is not required for timing of its inactivation in late mitosis.

Instead we find that FZR1 suppresses AURKA-dependent activity of the FRET biosensor in interphase. In the absence of FZR1 we speculate that the timing of reactivation of AURKA depends on the changing balance of AURKA activators and phosphatases during G1 phase. Although a low level of pT288 is detectable at centrosomes by IF in FZR1^KO^ cells, since there are routes to generating active AURKA that do not depend on autophosphorylation (for example under interaction with Nucleophosmin/B23 or TACC3 [46, 47], it is likely that basal AURKA activity in interphase is dependent on AURKA levels in the cell, and is calibrated in each cell cycle by mitotic destruction of the AURKA pool. Such multiple ‘opportunities’ for AURKA activation may underlie the strong association of AURKA overexpression with cancer, particularly where co-overexpressed with an activating partner such as TPX2 [48], and underscores the importance of ubiquitin-mediated destruction of AURKA in suppressing excess AURKA activity in interphase.

We investigated further the route to inactivation of the mitotic pool of AURKA. We found as expected that pT288 dephosphorylation during mitotic exit could be triggered through loss of interaction with TPX2. TPX2 has been shown to protect AURKA from negative regulation by PP1 [15–17, 49]. Indeed, whereas PP6 has been shown to act on the AURKA-TPX2 complex to limit AURKA activity in spindle assembly, PP1 can act only on ‘free’ AURKA [50]. We found that overexpression of the TPX2 (1-43) peptide that is sufficient to activate AURKA, and which is not degraded at mitotic exit, delayed the timing of kinase inactivation.

Our data indicate that the ‘release’ of AURKA to PP1-mediated inactivation could depend on APC/C-CDC20-mediated destruction of TPX2. We cannot exclude that ubiquitination of TPX2 would be enough to abrogate interaction, or that loss of interaction would be an indirect consequence of APC/C-CDC20 activity. However, our idea is consistent with the early degradation of TPX2 that we observe, and studies indicating TPX2 as a major substrate for APC/C at mitotic exit [51, 52] and numerous observations that amplification of TPX2 is the rate-limiting event for increased AURKA activity associated with tumorigenesis [48, 53]. One interesting conclusion to be drawn from our model is that although AURKA destruction during mitotic exit is a FZR1-dependent event, AURKA inactivation during mitotic exit is dependent on CDC20, along with all of the other known critical events of mitotic exit. This helps to explain why mitotic exit occurs essentially normally in absence of FZR1. Indeed, FZR1 is not expressed during early embryonic cell cycles, and AURKA activity there is regulated by activation and inactivation, instead of periodic proteolysis. Throughout development AURKA destruction therefore remains ‘decoupled’ from mitosis in a manner that contrasts with most other substrates of APC/C-FZR1, that can also be targeted by APC/C-CDC20 [36]. We infer that the unusual relationship between APC/C and Aurora kinases preserves a pool of potential Aurora kinase activity with critical interphase functions, regulatable through proteostasis.

So, what are the FZR1-sensitive AURKA-dependent events of interphase? Previous studies have shown that AURKA activity acts as an upstream regulator of mitochondrial morphology through RALA-dependent and -independent pathways [11, 13]. Mitochondria are highly dynamic organelles that undergo constant fission and fusion to modulate connectivity of the mitochondrial network, allowing the cell to respond to metabolic demands and maintain a healthy mitochondrial network. Increased rates of fission occur in preparation for mitosis so that mitochondrial fragments can be equally distributed between daughter cells. Here we show that the presence of non-degraded AURKA delays reassembly of the mitochondrial network after cell division. Whilst there is increasing evidence that the dynamic state of mitochondria contributes to the overall metabolic state of the cell, our understanding of how this impacts progression in the cell cycle remains limited [44, 45]. Our study provides new evidence that APC/C-FZR1 control of AURKA activity in interphase is a critical parameter in regulation of mitochondrial dynamics.

## Materials and Methods

### Generation of U2OS FZR1^−/−^ knockout cell line (FZR1^KO^)

Two guide RNAs (GACGTCCGATTGGAACGGCG and GCCCTGCCTCGCCATGGACC) targeting the first exon of FZR1 were cloned into AIO-GFP nickase vector, a gift from Steve Jackson (Addgene plasmid # 74119) [54].

U2OS cells were transfected with 2 μg of plasmid by electroporation using the Neon Transfection System according to the manufacturer’s instructions (Thermo Fisher Scientific). 48 hours after transfection cells were sorted based on GFP into 96-well plates at a single-cell-per-well density for clonal expansion. Single-cell clones were validated for loss of FZR1 by sequencing the genomic locus, immunoblotting and live-cell imaging of APC/C substrates.

### Cell culture, synchronization and drug treatments

U2OS and FZR1^KO^ cells were cultured in DMEM (Thermo Fisher Scientific) supplemented with 10% FBS, 200 μM Glutamax-1, 100 U/ml penicillin, 100 μg/ml streptomycin, and 250 ng/ml fungizone at 37°C with 5% CO2. For mitotic exit synchronizations, cells were collected in mitosis by 12 h treatment with 10 μM STLC (Tocris Bioscience) to trigger the Spindle Assembly Checkpoint (SAC) and then released at different timepoints by treatment with 10 μM AZ3146 (Generon, Slough, UK), an inhibitor of the SAC kinase Mps1. Cells were synchronized at different cell cycle stages as follows: For G0, cells were starved for 48 h in DMEM without serum. For G1, G0-arrested cells were released into serum-containing media for 2 h. For G1/S, cells were incubated with media containing 2 mM thymidine for 16 h, washed with PBS, released into regular media for 12 h, and then incubated in media containing 2 mM thymidine for 15 h. S-phase cells were prepared by releasing G1/S phase cells into regular media minus thymidine for 5 h. For M-phase cell population, cells were incubated with 10 μM STLC for 12 h. Mitotic cells were then collected by shake-off.

Aurora kinase inhibitors MLN8237 (Stratech, Ely, UK), MK5108 (Axon Medchem, Groningen, Netherlands), ZM447439 (Generon) and AZD1152-HPQA (Sigma-Aldrich UK) were used at the doses indicated.

RPE-1 cells, and RPE-1 FRT/TO cell lines expressing AURKA-Venus and AURKA-nd-Venus were cultured as previously described [12].

### Plasmids and transient transfection

pVenus-N1-AURKA, AURKA-Δ32-66 [27, 28] and TPX2(1-43) [23] have been described in the publications cited.

The non-targeted AURKB FRET biosensor was a kind gift from Michael Lampson [41].

The AURKA-directed FRET biosensor was created as followed: We modified a FRET construct backbone containing mTurquoise and mVenus sequences in the pECFP-C1 Clontech vector (construct “F36” [55], a kind gift from Carsten Schultz). First, mVenus was replaced by YPet sequence using KpnI and BamHI restriction sites. The sequence of the FHA2 domain from ScRad53 was amplified by PCR and inserted between AgeI and MluI sites, introducing HindIII and EcoRI sites upstream of MluI. Finally, complementary oligonucleotides encoding the phosphorylation site Thr210 of Plk1 were annealed and inserted between EcoRI and MluI sites. Amino acid sequence used: NH_2_-*G-G-S-G-G*-K-V-Y-D-G-E-R-K-K-K-**T**-L-C-I-COOH. Note that an isoleucine was introduced at the +3 position for FHA2 binding. The final plasmid construct contains the following elements: NcoI-mTurquoise-BglII-AgeI-FHA2-HindIII-EcoRI-Plk1 T210 phosphorylation sequence-MluI-YPet-BamHI. Complete sequence is available upon request.

Cells were transfected using electroporation with Neon Transfection System (Invitrogen™) using the following parameters: pulse voltage 1500 V, pulse width 10 ms, and 2 pulses total on the transfection device according to the manufacturer’s protocol.

### Immunoblotting

Cells were lysed in in 1% Triton X-100, 150 mM NaCl, 10 mM Tris–HCl at pH 7.5 and EDTA-free protease inhibitor cocktail (Roche), and PhosSTOP™ inhibitor for phosphatase (Sigma-Aldrich). After 30 min on ice, the lysate was centrifuged at 14,000 rpm (4°C) for 10 min. For immunoblotting, an equal amount of protein (20 μg) were loaded into SDS-PAGE 4-12% pre-cast gradient gels. Proteins were transferred to Immobilon-P or Immobilon-FL membranes using the XCell IITM Blot Module according to the manufacturer’s instructions. Membranes were blocked in PBS, 0.1% Tween-20, 5% BSA and processed for immunoblotting. Primary antibodies for immunoblot were as follows: AURKA mouse mAb (1:1000; Clone 4/IAK1, BD Transduction Laboratories), phospho-Aurora A (Thr288)/Aurora B (Thr232)/Aurora C (1:1000; clone D13A11 XP® Rabbit mAb, Cell Signalling), rabbit polyclonal TPX2 antibody (1:1000; Novus Biological), Cdh1 mouse mAb (1:50; gift from T. Hunt and J. Gannon), CDC20 mouse mAb (1:1000; Santa Cruz sc13162), AURKB rabbit polyclonal antibody (1:1000; Abcam ab2254), mouse monoclonal Cyclin B1 (1:1000; BD 554177), DRP1 rabbit polyclonal (1:500; Bethyl lab), rabbit polyclonal Tubulin (1:2000; Abcam ab6046), mouse mAb anti-Vinculin (1:1000; clone hVIN-1, Sigma-Aldrich), rabbit anti-GFP (1:1000; 11814460001, Roche). Secondary antibodies used were HRP-conjugated, or IRDye® 680RD- or 800CW-conjugated at 1:10000 dilution for quantitative fluorescence measurements on an Odyssey® Fc Dual-Mode Imaging System (LICOR Biosciences). Quantitative immunoblotting was carried out using IRDye® 680RD and 800CW fluorescent secondary antibodies, scanned on an Odyssey® Imaging System (LI-COR Biosciences).

### Immunofluorescence analysis

Cells were seeded at 2 × 10^4^ onto glass coverslips and then fixed with cold 100% methanol (−20°C), permeabilized with 0.5% Triton X-100 in PBS, and incubated in 2% bovine serum albumin (BSA), 0.2% Triton X-100 in PBS (blocking buffer) for 1 h at room temperature. Cells were incubated overnight with primary antibodies (rabbit phospho-AURKA Thr288, 1:50; mouse anti-γ-tubulin GTU-88, Sigma-Aldrich; 1:1000) diluted in blocking buffer at 4°C, then washed three times with blocking buffer and incubated with secondary antibodies at 1:1000 dilution. Alexa Fluor 488 anti-mouse and Alexa Fluor 568 anti-rabbit (Thermo Fisher Scientific) were used as the secondary antibodies. DNA was stained with DAPI. Coverslips were mounted with Prolong Gold antifade reagent. Epifluorescent stacks were acquired using 500 nm step with 2×2 bin using appropriate filter sets and 40X NA 1.3 oil objective. The best in-focus images were selected and integrated intensities were measured by ImageJ (http://rsb.info.nih.gov/ij/ National Institutes of Health, Bethesda, MD).

### Mitochondrial imaging and analysis

Cells were seeded at 2 × 10^4^ onto eight-well plastic-bottom slides (Ibidi GmbH, Martinsried, Germany) for live cell imaging. To stain the mitochondria, the cells were incubated with 100 nM Mitotracker^®^ Red CMXRos (M7512, Thermo Fisher Scientific) for 15 min, which was then replaced with L-15 medium supplemented with FBS. Epifluorescent images were acquired with 40X NA 1.3 oil objective on an Olympus IX81 motorized inverted microscope (Olympus Life Science, Southend-on-Sea, UK). The automated imaging platform included PE4000 LED illumination source (CoolLED, Andover, UK), Retiga R6 CCD camera (QImaging, Birmingham, UK), motorized stage (Prior Scientific, Cambridge, UK) and 37°C incubation chamber (Solent Scientific, Segensworth, UK), all controlled by Micro-Manager (Edelstein 2014). Images were collected with 2×2 bin applied, exported as tiff files and analysed using MicroP [12, 56].

### Timelapse imaging and FRET quantification

Cells were imaged in L-15 medium with 10% FBS at 37°C using an automated epifluorescence imaging platform composed of Olympus IX83 motorized inverted microscope, Spectra-X multi-channel LED widefield illuminator (Lumencor, Beaverton, OR, USA), Optospin filter wheel (Cairn Research, Faversham, UK), CoolSnap MYO CCD camera (Photometrics, Tuscon, AZ, USA), automated XY stage (ASI, Eugene, OR, USA) and climate chamber (Digital Pixel, Brighton, UK) and controlled using Micro-Manager. FRET imaging was performed using a 40X NA 0.95 objective and ECFP/EYFP/mCherry beamsplitter (Chroma, Bellows Falls, VT, USA) for ratiometric comparison of CFP and YFP emission upon excitation of CFP. ImageJ software (National Institutes of Health) was used to quantify CFP and YFP signal across the whole cell and AURKA activity expressed as CFP/YFP ratio (1/FRET).

### Statistical Analysis

Data analyses were performed in GraphPad 6.01 (San Diego, CA, USA). Results were analyzed with Student’s t-test or Mann-Whitney U-test (non-parametric) as indicated in figure legends. Significant results are indicated as p < 0.05 (*), p ≤ 0.01 (**) or p ≤ 0.001(***). Values are stated as the mean ± standard deviations.

## Acknowledgements

The authors thank Steve Jackson’s lab for the gift of AIO-GFP plasmid, Carsten Schultz for plasmid F36 and Michael Lampson for AURKB biosensor. We are grateful to Chiara Marcozzi for her advice on CRISPR/Cas9 and to Lilia Gheghiani for the design of FRET constructs. This work was funded by the MRC (MR/M01102X/1) and a Yousef Jameel Scholarship from the Cambridge International Trust to AA. OG is funded by Agence Nationale de la Recherche (ANR-15-CE13-0001) and GG by the Italian Association for Cancer Research (AIRC IG-17390). CL and GG acknowledge support from a Royal Society International Exchanges Award (IES/R3/170195).

## Author contributions

Project was conceived by CL, devised by CL, AA & HBA and carried out by AA whilst HBA created and characterized FZR1^KO^ cells and contributed experiments in Figure 2. MP carried out characterization of FRET biosensor (Figures 3 and S3). RG contributed experiments shown in Figures 6 and S5. Manuscript was written by AA & CL, results and manuscript discussed by all authors.

